# Investigating Bacterial Contributions to Thermal Tolerance in Three Intertidal Marine Snail *Tegula* Species

**DOI:** 10.1101/2025.05.13.653156

**Authors:** Brian Applegate, Meghan Burkhart, Hunter Caddow, Brighton Gover, Mary-Frances Kantola, Janessa Gaetos Obenchain, Anissa Smith, Bruce Nash, Ray A. Enke, Lani U. Gleason

## Abstract

In this era of climate change there is an urgent need to better understand the mechanisms that allow organisms to thrive vs. fail in thermally stressful environments. In particular, there is growing evidence that the “holobiont” (host animal + microbiome community of bacteria, fungi, and archaea that live in an organism) affects how organisms respond to environmental stressors such as temperature and thus should be studied further. Rocky intertidal species such as *Tegula* snails are ideal organisms for these types of studies because closely related species exhibit variability in heat tolerance. Here, we assess potential microbiome bacterial contributions to thermal tolerance in *Tegula eiseni, Tegula funebralis*, and *Tegula gallina* that co-occur in southern California but occupy different intertidal heights that vary in thermal stress exposure. 16S sequencing of the V4 region of individuals of each species exposed to control or 5.5-hour heat stress conditions (maximum temperature = 34 °C) revealed distinct bacterial communities across species and temperature treatments. Moreover, unique bacteria of the microbiome were significantly enriched in each *Tegula* species. For example, *Lutimonas, Polaribacter*, and the exopolysaccharide (EPS)-producing bacteria *Pelagicoccus* was most abundant in *T. gallina*, the species that occupies the highest intertidal heights and thus experiences heat stress most frequently. These results suggest that microbiome-derived metabolites such as EPS could be contributing to the higher thermal tolerance of *T. gallina*. Overall, this study demonstrates that the bacterial microbiome should be considered when examining mechanisms of thermal tolerance in marine invertebrates.

## Introduction

Climate change is on track to cause a sixth mass extinction event. Warming temperatures have thus far caused the extirpation of 400 species (IPCC, 2022), and one third of marine animals could become extinct in the next 300 years (Penn & Deutsch, 2022). Thus, there is an urgent need to better understand the mechanisms that differentiate organisms that thrive vs. fail in thermally stressful environments. In particular, there is growing recognition among organismal biologists that the “holobiont” (host animal + microbiome community of bacteria, fungi, and archaea that live in an organism) functions as an integrated unit (Lynch & Hsiao, 2019) and affects how organisms respond to environmental stressors such as temperature (Alberdi et al., 2016; Hector et al., 2022). Specifically, the microbiome is hypothesized to be associated with thermal tolerance in a diversity of organisms including lizards (Moeller et al., 2020), flies (Moghadam et al., 2017), aphids (Dunbar et al., 2007), corals (Ziegler et al., 2017), frogs (Fontaine et al., 2022), and several types of algae (Quigley et al., 2020; Xie et al., 2013).

Specific bacteria of the microbiome can affect thermal tolerance of the host by stimulating increased expression of stress response pathways genes, such as heat shock proteins (Brumin et al., 2011; Nakagawa et al., 2016; Porras et al., 2020) or by producing protective metabolites and proteins (Burke et al., 2010; Dunbar et al., 2007; Fontaine & Kohl, 2023). For example, some bacteria produce exopolysaccharides (EPS), complex sugar polymers excreted into the external environment that can protect against extreme stress conditions such as high temperatures, low nutrients, drought, salinity stress, and antimicrobial agents (Liu et al., 2013; Nichols et al., 2005; Wagh et al., 2022; Wang et al., 2019). There is evidence in tomato plants that EPS produced by plant-associated plant-growth promoting rhizobacteria reduce the negative effects of heat stress in the host plant (Morcillo & Manzanera, 2021). Whether EPS produced by the microbiome could also be playing a role in the thermal tolerance of marine invertebrate hosts remains unknown.

Rocky intertidal organisms are commonly studied when examining mechanisms of thermal tolerance because temperatures in the intertidal vary across small spatial scales and microhabitats, enabling comparisons of the same or related species that regularly experience different degrees of thermal stress. Previous work investigating microbiome differences in the intertidal has demonstrated that the host microbiome varies across these microhabitats and across temperature exposures. For example, 80% of the bacterial assemblage differs across bivalve clams *Ruditapes philippinarum* outplanted to three different intertidal levels that vary in emersion time (Offret et al., 2020). The cirri microbiome of the intertidal barnacle *Semibalanus balanoides* also varied across low vs. high intertidal microhabitats, and *Fucus* algae congeners occupying different heights of the intertidal zone vary in their microbiome composition and structure (Quigley et al., 2020). Similar differences have also been observed when comparing the microbiome of intertidal mollusks across temperature treatments: the bacterial community composition of the intertidal Sydney rock oyster *Saccostrea glomerata* changes following exposure to elevated heat wave temperatures (Scanes et al., 2023). Microbial communities also changed in response to elevated temperature in the mussel *Mytilus galloprovincialis* (Li, Xu, et al., 2019).

In this study we focus on the *Tegula* genus of intertidal marine snails on the west coast of the United States, looking specifically at *T. eiseni, T. funebralis*, and *T. gallina*. These species are well positioned to address questions regarding biological responses to climate change because they occupy overlapping but distinct geographic ranges and ecological niches (Hellberg, 1998). All three species co-occur in southern California. *T. eiseni* occupies the shallow subtidal zone (Schmitt, 1982) and is thus submerged underwater more and exposed to extreme high air temperatures less than the other two species. Conversely, *T. gallina* occupies the high intertidal zone, coinciding with more frequent and prolonged exposure to thermally stressful high air temperatures. *T. funebralis*, whose tidal height ranges from +0.4 to +2.0 m above mean lower low water (MLLW) in southern California (Gleason & Burton, 2016), occupies the mid to high intertidal zone. *T. funebralis* and *T. gallina* co-occur at roughly the same tidal heights in La Jolla and Bird Rock in San Diego County, California, but the range of *T. gallina* extends higher (authors’ unpubl. data). The phylogeny (Hellberg, 1998), heat shock response (Tomanek, 2002, 2005; Tomanek & Sanford, 2003; Tomanek & Somero, 1999, 2000), and transcriptome-wide response to heat stress (Gleason & Burton, 2015) of these species are well characterized, and the field microbiomes of *T. eiseni* and *T. funebralis* in southern California have also been examined (Neu et al., 2019). However, to date no work has been done comparing the microbiome of these species exposed to varying temperatures. Thus, our knowledge about how host species *and* heat stress affect the *Tegula* microbiome remains limited.

The objective of this study was to examine whether the microbiome contributes to thermal tolerance in *Tegula* intertidal snail species that occupy different tidal heights and thus experience distinct levels of thermal stress in the field. We exposed individuals of each *Tegula* species (*T. eiseni, T. funebralis*, and *T. gallina*) to control vs. heat stress conditions and performed 16S sequencing to characterize the bacterial community present in each condition. Specifically, we used the microbiome datasets to address the following questions: (1) does the bacterial community differ across *Tegula* species?; (2) does the bacterial community differ across control vs. heat stress treatments?; and (3) which specific bacteria are enriched in thermally tolerant and thermally sensitive *Tegula* species?

## Methods

### Microbiome Sample Preparation

Medium sized *Tegula eiseni, Tegula funebralis*, and *Tegula gallina* adults 15-20mm in shell diameter were collected in the summer of 2022 from the southern California site Bird Rock in San Diego County (32°48’N, 117°15’W). Snails were transported to California State University, Sacramento (Sac State) within 12 hours of collection. At Sac State snails of all three species were kept in a flow-through recirculating saltwater aquarium system set to 15 °C. Snails were regularly fed dried green algae sheets *ad libitum*. Before the start of any experiments, snails were kept at these temperature and food conditions for a common garden acclimation period of 3 weeks.

Following the common garden period *T. eiseni, T. funebralis*, and *T. gallina* individuals were exposed to either 1) control 15 °C conditions (*n = 3* per species) or 2) a 34 °C heat stress in air over 5.5 hours, following the same 3 °C increase per 30 min ramping protocol as described in Gleason and Burton 2013 (*n = 3* per species). At the conclusion of the heat stress, snails were frozen in liquid nitrogen and kept at -80 °C. The shell of each individual was removed with a woodworking vise, the remaining body was rinsed with ethanol, the gonads and muscular foot were removed, and DNA was extracted from 250 mg of the remaining whole-body tissue using a Qiagen DNeasy PowerSoil Pro kit. This tissue processing ensured that 1) microbiome differences based on gonadal sex differences did not influence the data, and that 2) the microbiome living in/on organs such as the lung, stomach, heart, etc. were examined (as opposed to the exterior surface bacteria on the muscular foot). A previous study found no differences in the microbiome of individual organs in *T. funebralis* and *T. gallina* (Neu et al 2019), and thus whole-body tissue samples were used here. All extracted DNA was quantified with a UV-Vis Nanodrop spectrophotometer (range 51.2 – 329.9 ng/uL).

### 16S Sequencing

A 350 bp region of the bacterial V4 region of the 16S rRNA gene was amplified using indexed forward (5’-GTGCCAGCMGCCGCGGTAA-3’) and reverse (5’-GGACTACHVGGGTWTCTAAT-3’) primers (Kozich et al., 2013) using the following protocol: 94 °C for 3 min, 35 cycles of 94 °C for 45 sec, 50 °C for 60 sec, 72 °C for 90 sec, and finally 72 °C for 10 min. PCR reactions were performed in 25 uL volumes and contained 12.5 uL 2x Phusion High-Fidelity DNA Polymerase Master Mix (NEB), 5 uL ddH2O, 1.25 uL of each primer (10 uM), and 5 uL of template DNA. Amplifications were verified on a 1% agarose gel with GelRed. A double-sided bead cleanup was carried out to remove primer-dimers and a low amount of off-target larger PCR products.

Quality and concentration of the pooled library was checked using a Bioanalyzer (Agilent, Santa Clara, CA, USA) and NEB’s Library Quant Kit for Illumina yielding 300bp PE reads. The library was then sequenced on an Illumina MiniSeq using a mid-output reagent cartridge at the James Madison University Center for Genome and Metagenome Studies (CGEMS). Before loading, the library was combined with Illumina’s PhiX control (30:70 16S:PhiX) to ensure a high-quality run despite the low diversity of the 16S library.

### Sequence Read Processing and Bacterial Identification

All 16S rRNA amplicon sequences were processed using the QIIME2 bioinformatics pipeline as implemented in the Purple Line of DNA Subway (Bolyen et al., 2019). DADA2 (Callahan et al., 2016) was used to trim low quality reads using the following parameters: truncLenF = 250; truncLenR = 231. After trimming to retain only high-quality reads, all samples were rarefied to a maximum depth of 33,529 sequences. We used a sampling depth of 3933 and the classifier Greengenes (515F/806R) to identify bacterial taxa in each sample.

### Assessment of Microbiome Differences

Alpha diversity was calculated in the Purple Line of DNA Subway using Pielou’s evenness (Pielou, 1966) and Faith’s phylogenetic diversity (Faith, 1992). We used Bray-Curtis distances to assess beta diversity of samples. NMDS plots were created using the *metaMDS* function in the R package *vegan* (Oksanen et al., 2001). To determine whether species or treatment significantly affected the microbiome, we conducted a Permutational multivariate analysis of variance (PERMANOVA) using the *adonis2* function in the R package *vegan* with 9,999 permutations. We used the *betadisper* functions of the *vegan* package to assess PERMANOVA results for heterogeneity of variance. The *plot_bar* function of the *phyloseq* package was used to visualize the relative abundances of the 25 most common bacterial genera with abundance counts of 400 and above. We used the linear discriminant analysis Effect Size (LEfSe) tool (Segata et al., 2011) as implemented in the R package *yingtools2* (Taur, 2016/2025) to identify specific microbial taxa that are found significantly more often in each *Tegula* species compared to the others. The R package *ggplot2* was used to generate and customize all figures. Lastly, for each treatment (control vs. heat stress), bacterial genera that are found 1) only in a single species, 2) in two of the three species, and 3) in all three species were identified and visualized in Venn diagrams using the online tool available at https://bioinformatics.psb.ugent.be/webtools/Venn/.

## Results

### Microbiome Comparison Across Species and Treatments

A total of 348,662 reads were sequenced across all samples, and 1406 different bacterial Genera were identified (Figure 1). The dominant taxa identified in *Tegula* samples was the phylum *Proteobacteria*. There were no significant differences in alpha diversity across the three *Tegula* species using either pielou’s eveness or faith’s pd indices (Kruskal-Wallis [pairwise], *p>0*.*05*; Figure 2). PERMANOVA results indicate that the microbiome communities were significantly different across the three *Tegula* species (df = 2, F = 2.758, *p* = 0.0016, and betadisper *p* = 0.392) and control vs. heat stress treatments (df = 1, F = 1.958, *p* = 0.0511, and betadisper *p* = 0.0568; Figure 3). Furthermore, the interaction between species and treatment was also significant (df = 2, F = 1.789, *p* = 0.0370).

**Figure 1.**
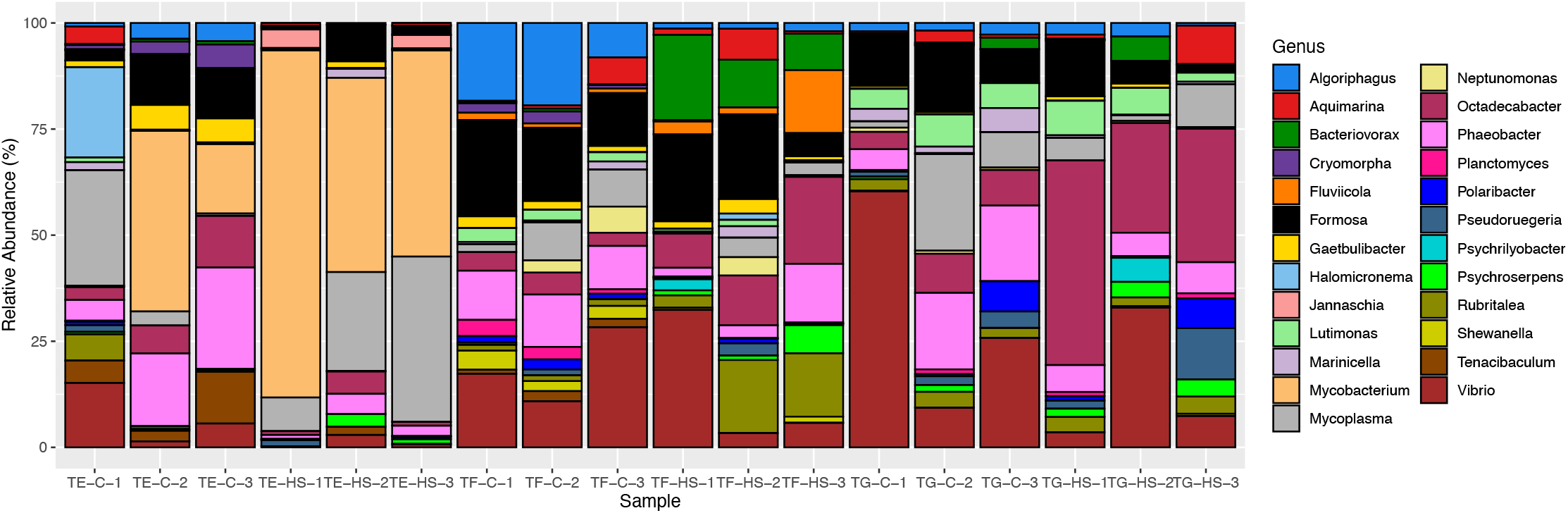
Taxa abundance plot of *Tegula* microbiomes under control (C) and heat stress (HS) conditions. Each vertical bar represents a different individual (TE = *T. eiseni;* TF = *T. funebralis;* TG = *T. gallina*), with bars grouped according to 1) species and 2) condition (*n* = 3). Each bar represents the relative abundance of bacterial taxa in that individual’s microbiome. Only taxa identifiable down to the Genus level and with more than 400 counts were included here.

**Figure 2.**
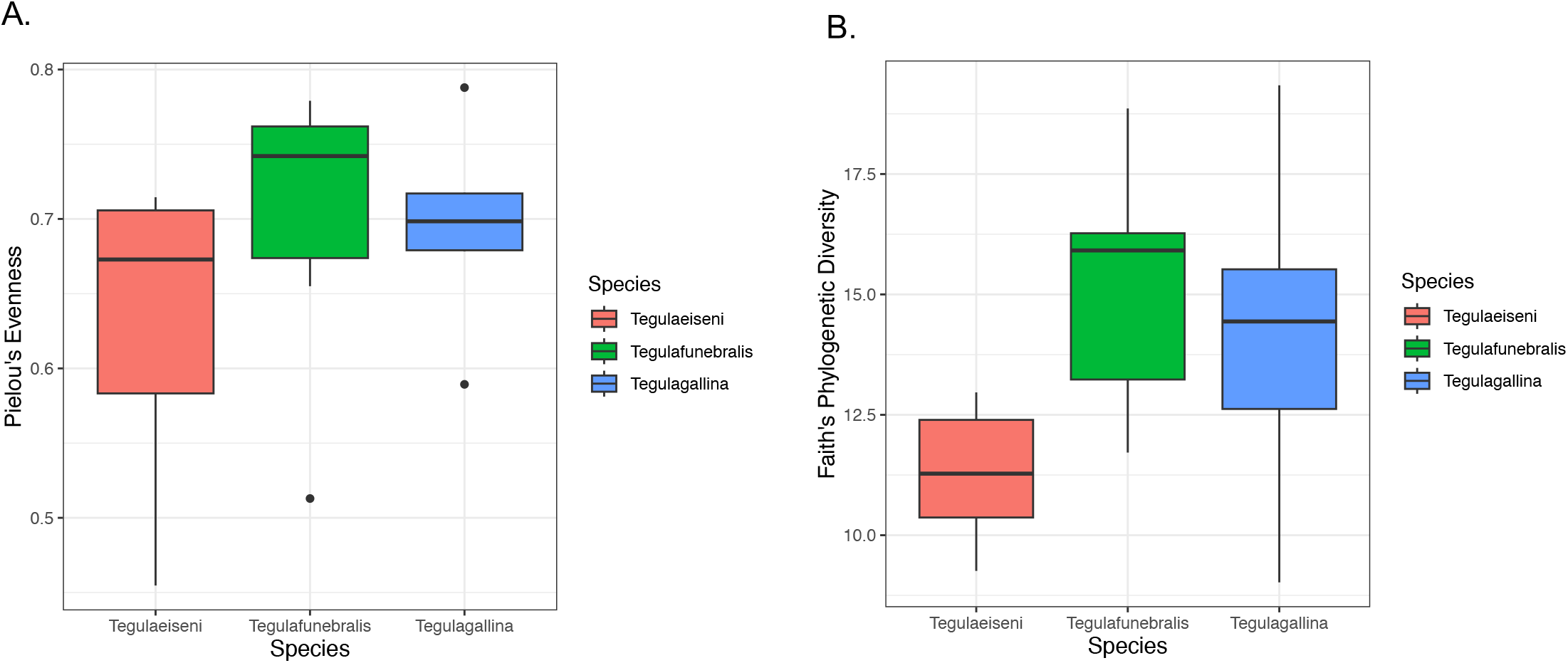
Alpha diversity across the three *Tegula* species calculated using Pielou’s Evenness (a.) and Faith’s Phylogenetic Diversity (b.) metrics.

**Figure 3.**
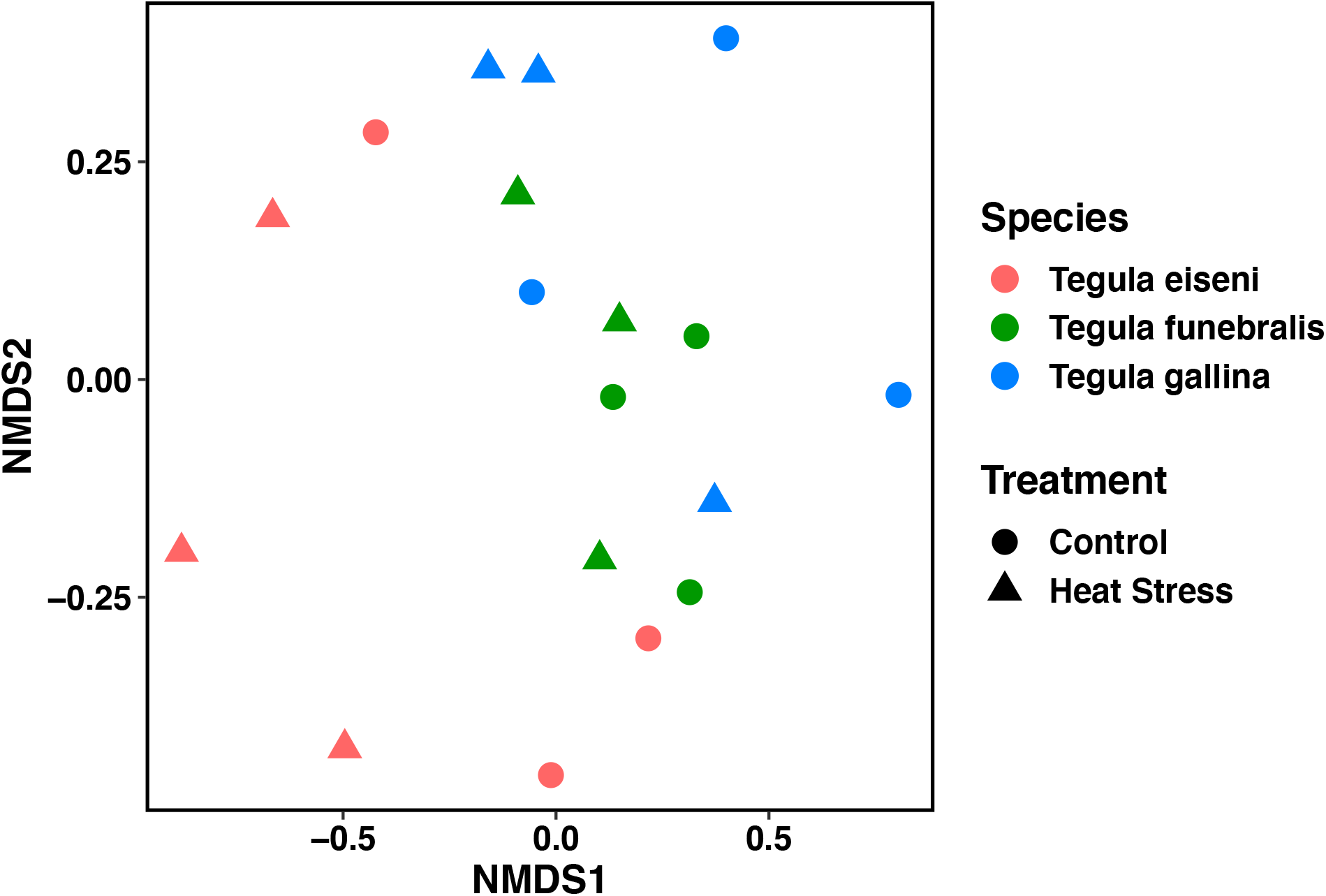
Non-metric multidimensional scaling (NMDS) plot of *Tegula* microbiomes under control (circles) and heat stress (triangles) conditions. *n = 3* for each treatment group (e.g., *T. eiseni* control). Each data point represents an individual snail, with *T. eiseni* shown in pink, *T. funebralis* in green, and *T. gallina* in blue.

### Enriched and Unique Microbial Taxa

Significantly enriched bacterial genera with an excess of reads were identified in each of the three *Tegula* species (Table 1). For *T. eiseni*, there was an excess of reads belonging to taxa in the genus *Tunicatimonas, Halomicronema*, and *Mycobacterium* (linear discriminant analysis [LDA] log scores = 2.89, 3.24, and 4.90, respectively; Figure 4). *T. funebralis* had four enriched bacterial genera: *Polynucleobacter, Shewanella, Fluviicola*, and *Rubritalea* (LDA log scores = 2.67, 3.59, 3.73, and 3.89, respectively). Lastly, the three bacterial genera *Pelagicoccus, Polaribacter*, and *Lutimonas* (LDA log scores = 2.12, 3.26, and 3.52, respectively) had an excess of reads in *T. gallina*.

**Table 1.**
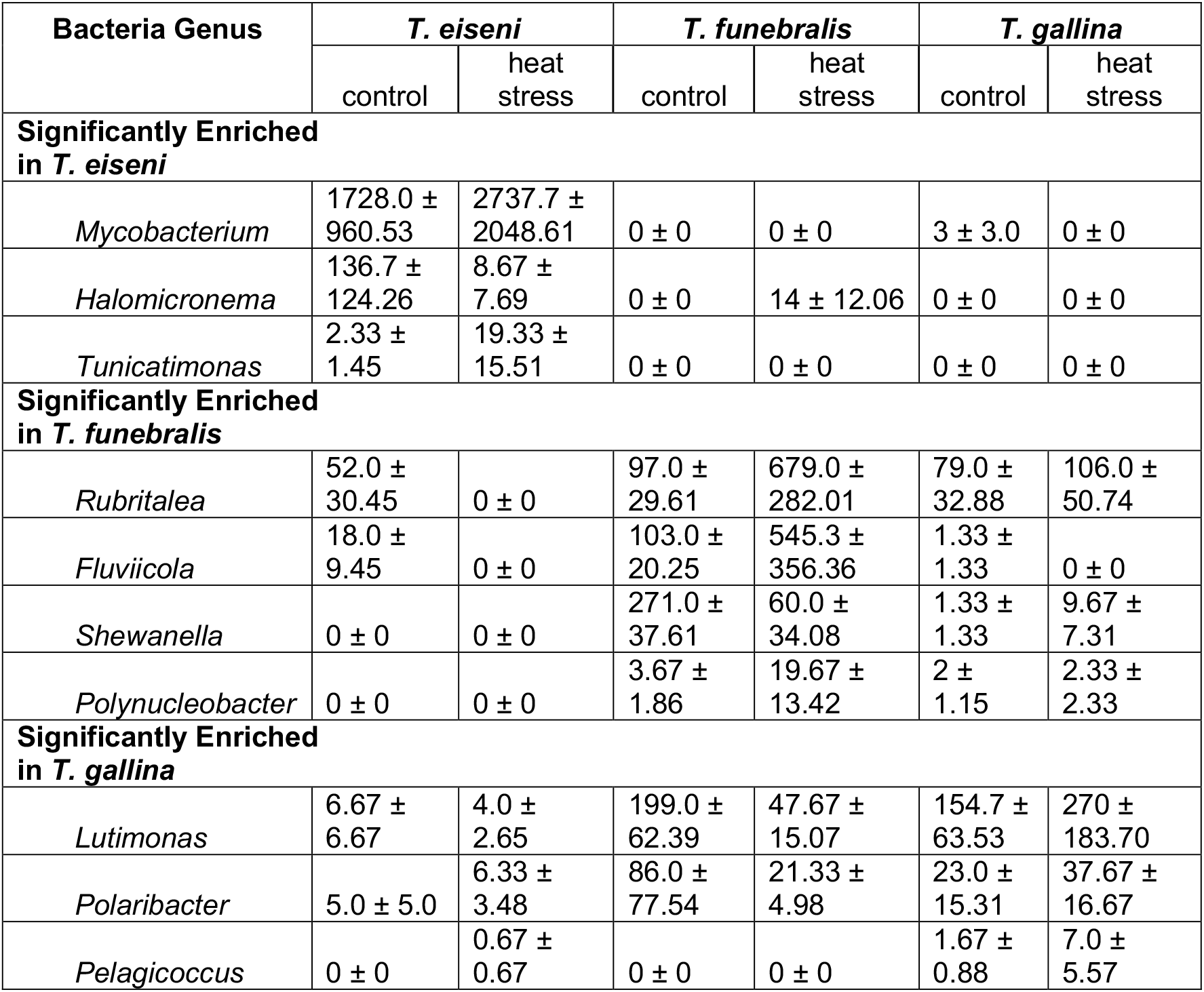
Average abundance in raw number of sequences (±SEM) of bacteria genera identified in linear discriminant analysis (LDA) to be significantly enriched in (i) *T. eiseni*, (ii) *T. funebralis*, and (iii) *T. gallina*. For each species, data for control and heat stress conditions are presented in separate columns.

**Figure 4.**
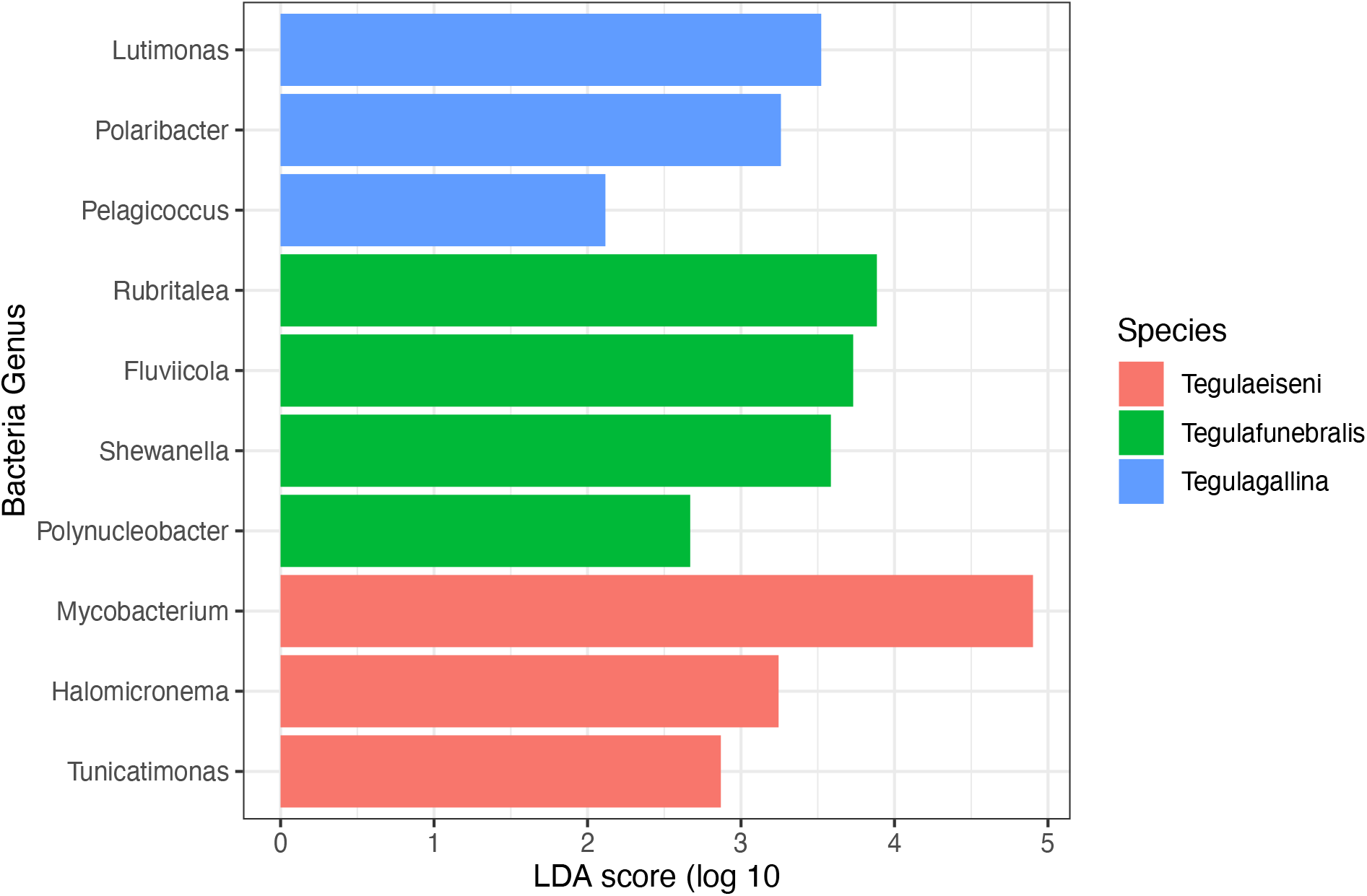
Linear discriminant analysis (LDA) log scores for the significantly enriched bacterial Genera in each *Tegula* species (*T. eiseni* in pink, *T. funebralis* in green, and *T. gallina* in blue).

As shown in Figure 5a, under control conditions *T. eiseni* had the least unique bacterial taxa (36 Genera, 26.5% of the total genera identified; Supplemental Table 1), including three members of the *Flammeovirgaceae* family. In contrast, *T. gallina* had the most unique bacterial taxa (50 Genera, 38.2% of the total genera identified). Unique taxa in *T. gallina* include two members of each of the following families: *Alcaligenaceae, Cytophagaceae, Peptostreptococcaceae*, and *Rhodobacteraceae*. Under heat stress conditions this pattern was different: *T. gallina* had the least unique bacterial taxa (20 genera, 15.4% of the total genera identified), including two members each of the *Flavobacteriaceae* and *Verrumicrobiaceae* families. Heat-stressed *T. funebralis* had the most unique bacterial taxa (61 Genera, 32.3% of the total Genera identified), including three members each of the families *Moraxellaceae* and *Pirellulaceae* (Figure 5b).

**Figure 5.**
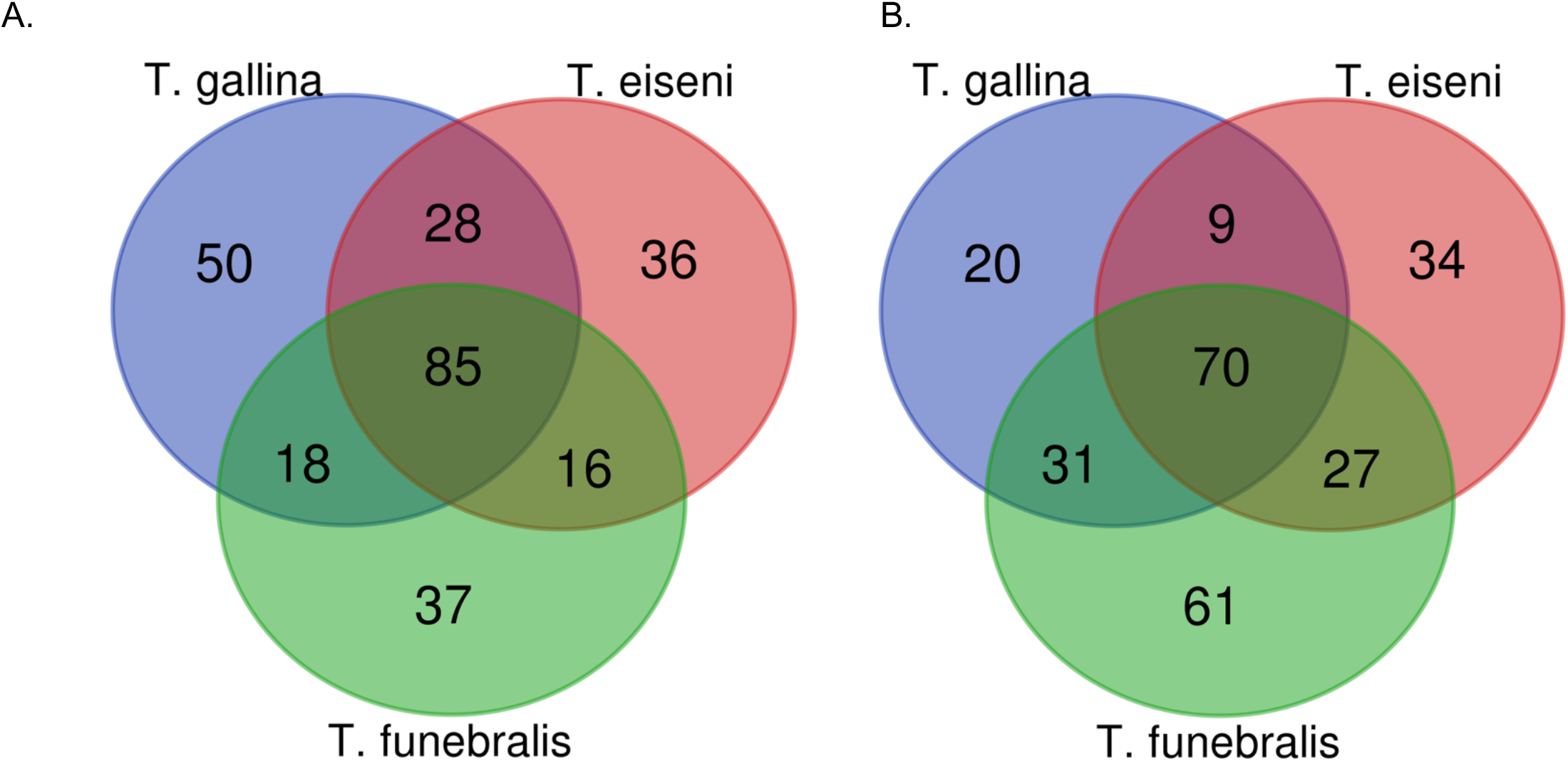
Number of unique vs. shared bacteria genera identified in each *Tegula* species under control (A.) and heat stress (B.) conditions.

There were three bacterial genera found only in *T. eiseni* under both control and heat stress conditions: *Owenweeksia, Tunicatomonas*, and *Jannaschia*. Five bacterial genera were only found in *T. funebralis* under both control and heat stress conditions: *Clostridium, Aquicella, Croninitomix, Plantomycete*, and *Roseovarius*. There were no taxa present only in *T. gallina* under both control and heat stress conditions.

## Discussion

The microbiome of the intertidal marine snails *T. eiseni, T. funebralis*, and *T. gallina* significantly differs across species and control vs. heat stress treatments. These differences are driven by both unique (i.e., bacteria that are only present in one of the three *Tegula* species) and enriched (i.e., bacteria that have a significant excess of reads in one of the three *Tegula* species) bacterial genera.

### Microbiome Communities Significantly Differ Across Species

Proteobacteria is the most common bacterial phyla across all three *Tegula* species – this is consistent with previous work in marine animals such as snails, limpets, red abalone, corals, copepods, fish, and barnacles (Bayer et al., 2013; Brown et al., 2020; Dorosz et al., 2016; Dudek et al., 2014; Givens et al., 2015; Neu et al., 2019; Ousley, 2023). However, at the genus level the microbial community composition significantly varies across *Tegula* species. This is consistent with previous work in *Tegula* (*Chlorostoma*) *eiseni* and *Tegula* (*Chlorostoma*) *funebralis* examined directly from the field from La Jolla, California (Neu et al., 2019). Our results also match findings in other marine species such as sponges and algae. In previous work in four different demosponge species, samples clustered together in non-metric multidimensional scaling (NMDS) space according to species (De Castro-Fernández et al., 2023). Moreover, in a study design similar to this one, the microbiome composition and structure significantly varied across three *Fucus* algae congeners occupying different heights of the intertidal zone and thus experiencing different levels of abiotic stress (Quigley et al., 2020).

Notably, our results provide evidence for phylosymbiosis, in which the microbiomes of more closely related species are more similar to each other. As seen in Figure 3, in multivariate NMDS space the microbiome of *T. eiseni* is distinct from the other two species *T. funebralis* and *T. gallina*, who are more closely related to each other. Such phylosymbiosis has also been observed when comparing the microbiomes of *Chlorostoma* (now *Tegula)* and *Littorina* intertidal snails (Neu et al., 2019) and of different tropical sponge species (Easson & Thacker, 2014).

Evolutionary codivergence of the microbiome with these *Tegula* species is one possible explanation for microbiome differentiation, but several other explanations, such as habitat filtering, should also be considered and investigated further (Mazel et al., 2018). Microbial communities are sensitive to environmental parameters such as pH and temperature; potential differences in the gut environment between *T. eiseni, T. funebralis*, and *T. gallina* could be contributing to their distinct microbiomes (Neu et al., 2019). Although our experiments were conducted in a common garden laboratory environment in which all species were fed the same dried algae food, differences could also be artifacts of the unique field microhabitats each species was collected from in the intertidal. For example, when discussing the microbiome differences between field-collected *T. eiseni* and *T. funebralis*, Neu et al hypothesized that these species could be differentially ingesting microbes through dietary inputs (Neu et al., 2019).

Moreover, the cirri microbiome differences in barnacles occupying low vs. high intertidal microhabitats was thought to be due to differential exposure to and/or time underwater (Brown et al., 2020). Comparing the microbiome of *T. eiseni, T. funebralis*, and *T. gallina* individuals collected directly from the field to those that have been in a lab common garden environment could help determine how much influence the distinct microhabitats of each species have on their microbiome. Similarly, explicitly manipulating diet and/or time underwater in a laboratory setting could also provide further insight into the mechanisms of microbial differentiation across species.

### Microbiome Communities Significantly Differ Across Treatments

In addition to variation across species, we also observed microbiome differentiation between individuals exposed to control vs. heat stress conditions. Such differentiation has also been observed in other intertidal marine mollusks. In *Haliotis rufescens* red abalone, another type of marine snail, control individuals had a distinct microbial composition compared to heat stressed individuals (Ousley, 2023). Similarly, in the intertidal Sydney rock oyster *Saccostrea glomerata* exposure to elevated heat wave temperatures significantly changed bacterial community composition (Scanes et al., 2023). Microbial communities also changed in response to elevated temperature in the mussel *Mytilus galloprovincialis* (Li, Xu, et al., 2019). One potential mechanism causing this microbial shift under higher temperatures is the proliferation of opportunistic pathogens. For example, one such pathogen *Mycobacterium* was found to be enriched in the heat sensitive *T. eiseni* as described below. Similar increases of pathogenic bacteria following exposure to heat stress have also been observed in seaweed (Düsedau et al., 2023), mussels (Li, Chen, et al., 2019), and oysters (Green et al., 2019).

### Enriched Bacteria in Heat Sensitive Species

In *T. eiseni*, the species with the lowest thermal tolerance, enriched bacteria could be contributing to infection and apoptotic cell death under heat stress conditions. *Mycobacterium* is much more abundant in *T. eiseni* under both control and heat stress conditions compared to *T. funebralis* and *T. gallina*. This is a pathogenic bacteria that is known to infect other marine mollusks (Davidovich et al., 2020), and enrichment of this taxa could suggest an increased risk of disease in *T. eiseni*, especially under heat stress conditions. Another taxa enriched in *T. eiseni, Halomicronema*, is a cyanobacteria that has also been found in marine sponges (Caroppo et al., 2012). Notably, some compounds produced by the *Halomicronema* genus can trigger apoptotic cell death (Mutalipassi et al., 2019). In other marine mollusks such as the oyster *Crassostrea virginica*, the density of apoptotic cells increased after exposure to high temperatures of 26 and 30 °C (Rahman & Rahman, 2021). These authors hypothesized that a high level of reactive oxygen species (ROS) could be contributing to this increase in apoptosis (S. Nash et al., 2019; S. Nash & Rahman, 2019). Our results suggest components of the microbiome could also be contributing to the induction of apoptosis in heat sensitive marine mollusks, and should be investigated further. The last enriched bacterial genus in *T. eiseni, Tunicatomonas*, has also been isolated from sea anemones, but there is only one known species, and not much information is available on this taxon (Yoon et al., 2012).

### Enriched Bacteria in Heat Tolerant Species

*Lutimonas* was found in relatively high abundance in *T. gallina* under both control and heat stress conditions. These results contrast a previous study that observed a decrease in *Lutimonas* under heat stress conditions in Pacific white shrimp *Litopenaeus vannamei* (Duan et al., 2021). *Lutimonas* is a nitrifying bacteria that degrades ammonium (Fu et al., 2009). In other marine mollusks such as the ark shell *Scapharca subcrenata* (Jiang et al., 2020) and in *Daphnia* (N. Nash et al., 2022), elevated temperatures increase ammonia excretion rates. High levels of ammonia are known to have an array of negative consequences for aquatic invertebrates (Zhang et al., 2023); thus, the relatively high abundance of *Lutimonas* in *T. gallina*, even under heat stress conditions, could suggest that the microbiome helps regulate ammonia levels to prevent toxic overaccumulation.

Our finding that the genus *Polaribacter* is enriched in *T. gallina*, including under heat stress conditions, contrasts a previous study in another marine mollusk *Mytilus galloprovincialis* that found *Polaribacter* was a dominant genus under control conditions, but its abundance decreased when water temperature was increased (Li, Xu, et al., 2019). However, in other marine invertebrates, including the mussel *Mytilus coruscus* (Li, Chen, et al., 2019) and the common yellow sponge *M. acerate* (De Castro-Fernández et al., 2023), abundance of *Polaribacter* was also high under heat stress conditions. *Polaribacter* has also been detected in sea water (Fukui et al., 2013; J.-H. Yoon et al., 2006) and in diatom phytoplankton blooms (Xing et al., 2015). Overall, not much is known about this genus, and further research must be done before hypotheses regarding the ability of this genus to increase heat tolerance of *T. gallina* can be formed.

The third and final bacterial genus that was enriched in *T. gallina* is *Pelagicoccus. Pelagicoccus*, a member of the family *Puniceicoccaceae*, contains four different species (Feng et al., 2021; J. Yoon, Oku, et al., 2007a). These species have previously been isolated from sea grass (J. Yoon, Oku, et al., 2007b), seawater (J. Yoon, Yasumoto-Hirose, et al., 2007), and marine sediment (Feng et al., 2021). Based on the genomic analysis performed by Feng et al, it is thought that all bacteria in this genus produce exopolysaccharides (EPS) to better cope with extreme stress conditions, including high temperatures, low nutrients, drought, salinity stress, and antimicrobial agents (Liu et al., 2013; Nichols et al., 2005; Wagh et al., 2022; Wang et al., 2019). For example, EPS are produced in hydrothermal vent communities exposed to extreme high temperatures (Nichols et al., 2005). Importantly, there is evidence in other species that EPS produced by bacteria have positive effects on their host: EPS produced by plant-associated plant-growth promoting rhizobacteria reduce the negative effects of heat shock and growth in plant hosts such as tomatoes (Morcillo & Manzanera, 2021). It is possible the *Pelagicoccus* observed in this study could similarly be contributing to host thermal tolerance in *T. gallina*, although it is not currently known if the relatively low abundance of *Pelagicoccus* would be sufficient to produce beneficial effects.

### Caveats

As with any study, there are limitations that should be considered. Most notably, we did not collect or sequence environmental samples to characterize the surrounding microbial habitat (e.g. seawater) of the *Tegula* snails. Although we did thoroughly rinse tissue samples before extracting DNA (Brown et al., 2020; Neu et al., 2019), we nevertheless cannot determine whether the *Tegula* microbiome communities we characterized are subsets of the surrounding bacterial community in the seawater and/or the algal diet of the snails. Moreover, at this point it is unknown if the bacterial species we observed in the *T. eiseni, T. funebralis*, and *T. gallina* microbiomes are transient or resident. To date, we have only characterized the microbiome following a short 5.5-hour heat stress representative of a low tide period in the field. No assessment has yet been performed of the recovery period following this heat stress; in other words, it is not yet known if the microbial community reverts to the same “baseline” composition after return to non-stress control conditions. In addition, this work investigates the microbiome from a single field site in San Diego, California. Further work is needed to determine if there are microbiome differences within a single *Tegula* species across different geographic regions.

## Conclusion

This study demonstrates that the microbiome of *Tegula* species is distinct under both control and heat stress conditions and suggests that components of the microbiome could be contributing to the differential thermal tolerance of *T. eiseni, T. funebralis*, and *T. gallina* intertidal snails. Specifically, the pathogenic bacteria *Mycobacterium* is significantly enriched in the thermally sensitive *T. eiseni*, and the nitrifying bacteria *Lutimonas* and the exopolysaccharide-producing bacteria *Pelagicoccus* are significantly enriched in the thermally tolerant *T. gallina*. Overall, our results provide further insight into how the microbiome differs across host congeners living in uniquely stressful microhabitats and illustrate the additional information gained when considering non-genetic mechanisms of thermal tolerance.

## Acknowledgements

We thank Caity Fox, Instructional Support Technician II at CSU Sacramento, for preparing reagents for sample processing and DNA extractions and Karen Barnard-Kubow at the JMU CGEMS for technical assistance with library preparation and DNA sequencing.

## Authorship Contribution Statement

BA, MB, HC, BG, M-FK, JGO, AS: formal analysis, investigation; RAE: investigation, resources, funding acquisition; BN: resources, funding acquisition; LUG: conceptualization, data curation, formal analysis, investigation, writing

## Supplemental Materials

Supplemental Table 1: Lists of bacteria found to be common or unique across the three *Tegula* species under control conditions, as shown in the Venn diagram in Figure 5a. Bacteria representing each region of the Venn diagram are listed in separate tabs (TE = *T. eiseni*, TF = *T. funebralis*, TG = *T. gallina)*. Each bacterium is listed across a single row, with the different taxonomic classifications (e.g. Kingdom, Phylum, Class, Order, Family, Genus) shown in separate columns.

Supplemental Table 2: Lists of bacteria found to be common or unique across the three *Tegula* species under heat stress conditions, as shown in the Venn diagram in Figure 5a. Bacteria representing each region of the Venn diagram are listed in separate tabs (TE = *T. eiseni*, TF = *T. funebralis*, TG = *T. gallina)*. Each bacterium is listed across a single row, with the different taxonomic classifications (e.g. Kingdom, Phylum, Class, Order, Family, Genus) shown in separate columns.

